# Encoded metabolic remodeling amplifies drug resistance in *Mycobacterium tuberculosis*

**DOI:** 10.64898/2026.05.05.722911

**Authors:** Abigail M Frey, Gregory H. Babunovic, Peter H. Culviner, Xin Wang, Ella Meirav, Mingyu Gan, Junhao Zhu, D Branch Moody, Qingyun Liu, Sarah M. Fortune

## Abstract

Antibiotic pressure causes pathogens to evolve many forms of altered drug susceptibility. In addition to target or activator mutations conferring canonical drug resistance, mutations can serve as steppingstones to or enhancers of resistance. In clinical strains of *Mycobacterium tuberculosis* (Mtb), we find that *idsA2,* which encodes an isoprenyl pyrophosphate synthase involved in the synthesis of precursors for essential components of the cell wall and electron transport chain, is undergoing diversifying selection and that these mutations are associated with the acquisition of first-line antibiotic resistance. By engineering isogenic Mtb strains that express clinically prevalent variants of *idsA2*, we show that clinical variants increase the minimum inhibitory concentration of ethambutol and, to a lesser extent, of isoniazid. Targeted lipid analyses reveal that disrupting IdsA2 function redirects limited resources in the isoprenoid synthesis pathway, leading to increased production of decaprenylphosporyl pentose which can compete with ethambutol for binding to arabinosyltransferases. *IdsA2* mutations most often occur after *embB* mutation and lead to a multiplicative increase in ethambutol resistance. Thus, identification of *idsA2* mutations can be utilized to improve the specificity of genotypic ethambutol susceptibility testing. Together, this work defines *idsA2* as an ethambutol resistance gene and demonstrates how metabolic remodeling can augment drug resistance.

**Author Summary:** Tuberculosis is the deadliest infectious disease in the world and becomes more difficult to treat with the acquisition of drug resistance. Defining the mechanisms and genetic basis of drug resistance will allow for improved screening and drug regimen optimization for treatment success. To identify previously unrecognized mechanisms of altered drug susceptibility, we have combined population genomics and experimental genetics approaches, focusing on *Mycobacterium tuberculosis* genes evolving in clinical strains. We identified *idsA2* as a target of frequent mutations that are associated with drug resistance. Experimental data indicate that *idsA2* variants decrease the susceptibility of the bacteria to multiple antibiotics through isoprenoid synthesis remodeling, with the strongest effect on ethambutol resistance. *IdsA2* mutations often occur after *embB* mutations to multiplicatively increase ethambutol resistance. These data suggest that inclusion of *idsA2* variants in drug resistance testing could improve the specificity of genotypic detection of ethambutol resistance.

## Introduction

While traditionally thought of as monomorphic, the importance of the genetic variation of *Mycobacterium tuberculosis* (Mtb) has become increasingly evident. This variation has evolved over both short and long timescales, resulting in heterogeneity within a single patient as well as distinct phylogenetic lineages of Mtb across the globe [1]. Drug resistance mutations are some of the most prevalent variants in Mtb [2,3], illustrating the strength of antibiotics as a selective pressure on the Mtb genome. Although the best understood mutations selected by antibiotic pressure cause high-level drug resistance, drug pressure also selects for other classes of mutations that give Mtb a fitness advantage during treatment. Drug tolerance mutations allow Mtb to survive longer in the face of drug stress [4]; resilience mutations shorten the period of post-antibiotic growth arrest [2]; and compensatory mutations mitigate the fitness defects caused by high-level resistance mutations [5]. Furthermore, while resistance mutations are typically found in drug activators or targets [6], low-level resistance mutations that indirectly but meaningfully alter antibiotic action have been associated with treatment failure and enable acquisition of additional drug resistance variants [7,8].

The complexity of bacterial evolution under drug pressure can be effectively dissected by coupling population genomic and experimental approaches. Here we leverage this strategy to identify the impact of clinically prevalent mutations in *idsA2*, a common target of diversifying selection in the Mtb genome with variants that have been associated with drug resistance [9,10]. *IdsA2* encodes an isoprenyl pyrophosphate synthase (IdsA2) which catalyzes the synthesis of precursors used for essential processes such as building the cell wall and bacterial energetics [11]. As determined by transposon insertion screening, *idsA2* itself is predicted to be nonessential *in vitro*, putatively due to enzymatic redundancy [12–14]. Isoprenoid synthesis itself is not targeted by any antimycobacterial agents.

In this work, we leverage our Mtb variant dataset derived from over 55,000 clinical strains to show that *idsA2* is mutating for loss of function specifically in Lineage 4 Mtb strains, and these variants are associated with drug resistance. When engineered into Mtb*, idsA2* variants increase bacterial survival in the face of treatment with both ethambutol and isoniazid as a result of different consequences of isoprenoid synthesis pathway remodeling. Our phylogenomic and experimental analyses suggest that the effect on ethambutol susceptibility is clinically most relevant, where *idsA2* mutations combine with *embB* mutations to multiplicatively contribute to increased ethambutol resistance beyond critical concentrations [15]. The second-step nature of *idsA2* mutations could allow them to be utilized clinically on top of *embB* variants to improve the specificity of ethambutol resistance predictions. In total, we present *idsA2* variants as previously unappreciated genetic contributors to first-line drug resistance discovered through large-scale study of Mtb evolution.

## Results

### *idsA2* is under selection for loss of function

Antibiotics are a profound selective force on the Mtb population and many known drug resistance genes are under diversifying selection even over short time frames in a single individual. We previously identified *idsA2* as a frequent target of within-host selection [2]. Here we extended this work to determine if *idsA2* variants are under diversifying selection at the population scale, leveraging our variant dataset derived from the whole genome sequencing data of ∼55,000 clinical isolates (S1 Table); we refer to this dataset going forward as the 55K dataset [3]. Across all strains, the ratio of nonsynonymous to synonymous mutational events (*pN/pS*) in *idsA2* was 0.99; this was in the top 6% of *pN/pS* values across the Mtb genome (Fig 1A, S1 Table). While a *pN/pS* of 1 is usually interpreted as evidence of neutral selection [16], it could also reflect heterogeneous selective pressures across different lineages, which have different mutational backgrounds (S2 Table). Indeed, in Lineage 4, the *pN/pS* of *idsA2* was 1.29 which was within the top 3% of *pN/pS* values across the genome (Fig 1A, S1 Table). In Lineages 1, 2, and 3, the *pN/pS* value were closer to the median (S1 Fig), suggesting that the benefit or ability of *idsA2* to diversify is unique to Lineage 4.

**Fig 1:**
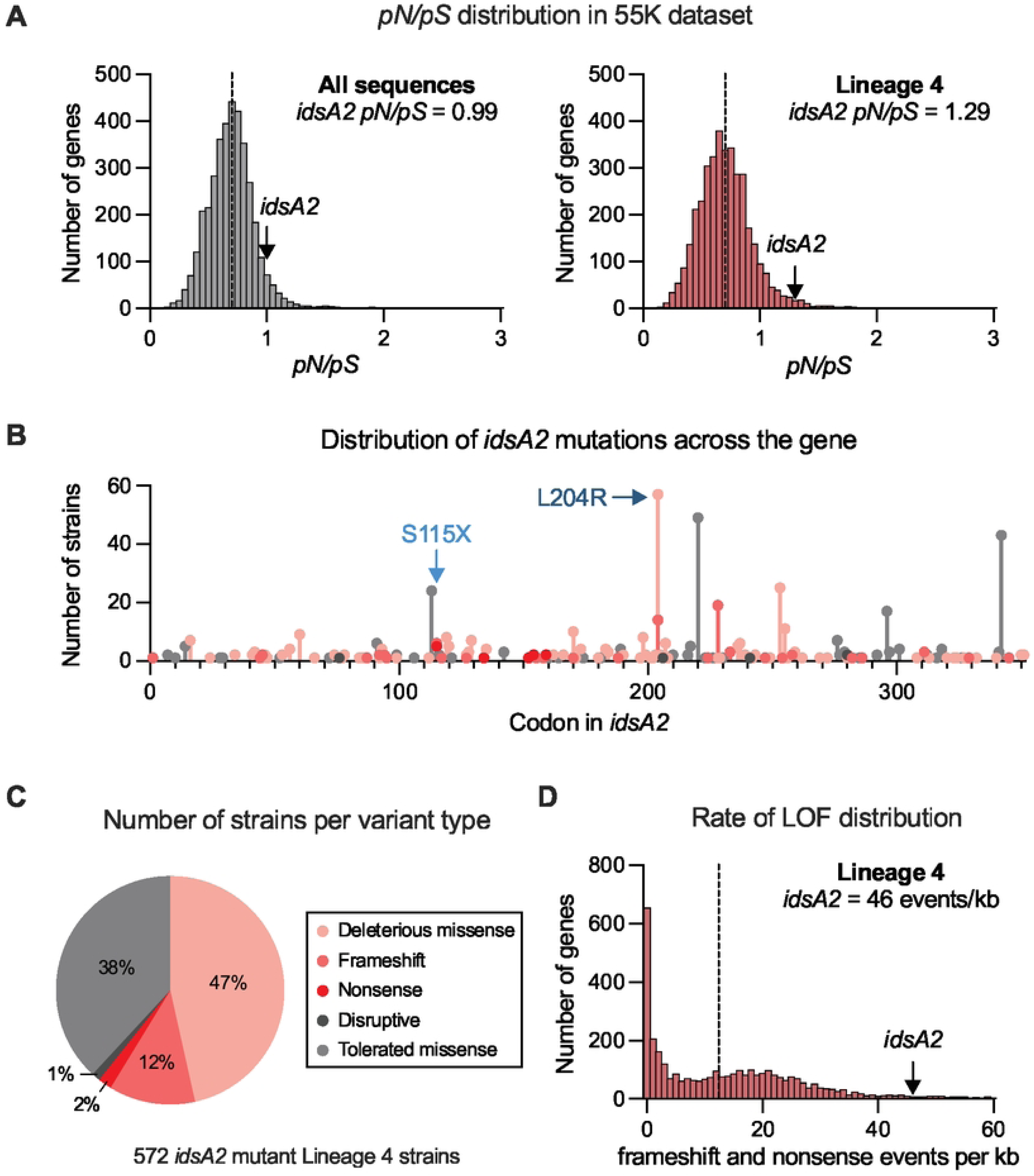
*idsA2* is under selection for loss of function. (A) Histograms showing the number of genes per *pN/pS* ratio across the 55K dataset (left) and Lineage 4 (right). Median values indicated by the dotted lines are both 0.69 and *idsA2 pN/pS* values indicated by the arrows are 0.99 and 1.29. (B) The number of strains per codon from Lineage 4 of the 55K dataset with disruptive in-frame deletions or insertions, frameshift, deleterious or tolerated missense, or nonsense mutations in *idsA2*. Strains with mutations introduced into H37Rv by recombineering are indicated. (C) Pie chart of the number of strains per *idsA2* variant type from Lineage 4 in the 55K dataset. (D) Histogram showing the number of genes per rate of frameshifts and nonsense variants across Lineage 4 of the 55K dataset. The median value indicated by the dotted line is 12.7 mutations/kb and the number of frameshifts and nonsense mutations in *idsA2* is 46.3 mutations/kb.

We next sought to predict the effect of *idsA2* mutations. In Lineage 4 strains, *idsA2* mutations occurred across the gene body, and 61% of strains were predicted to have a deleterious variant in *idsA2,* annotated as a frameshift, nonsense, or SIFT-predicted deleterious missense mutation (Figs 1B-C) [17]. To contextualize this genetic signal of loss of function, we compared the burden of nonsense and frameshift mutations in *idsA2* to that of other genes in the Mtb genome (S3 Table). *IdsA2* was in the top 6% of genes for frequency of frameshift and nonsense mutations (Fig 1D), suggesting that in Lineage 4, *idsA2* is under selection for loss of function mutations.

### In clinical strains, *idsA2* mutations are associated with drug resistance

We then assessed the association between *idsA2* mutations and antibiotic sensitivity using data from the CRyPTIC consortium which performed whole genome sequencing and measured minimum inhibitory concentrations (MICs) for first- and second-line drugs across over 10,000 Mtb strains (S2 Fig) [18]. We calculated the average difference in MICs between Lineage 4 strains with *idsA2* mutations and their phylogenetic nearest neighbor(s), assessing the significance of this difference through permutation testing. *IdsA2* mutational events were significantly associated with increases in the MICs of rifampicin, isoniazid, and ethambutol (Fig 2 and S3). The average difference in MIC was further increased when limited to only predicted-deleterious *idsA2* mutational events (Fig S4). Due to the concurrent use of these drugs for treatment of tuberculosis, mutations altering drug susceptibility are genetically linked, therefore, these data do not reveal which drug selects for *idsA2* mutations. Thus, we next sought to experimentally define the effect of *idsA2* mutations.

**Fig 2:**
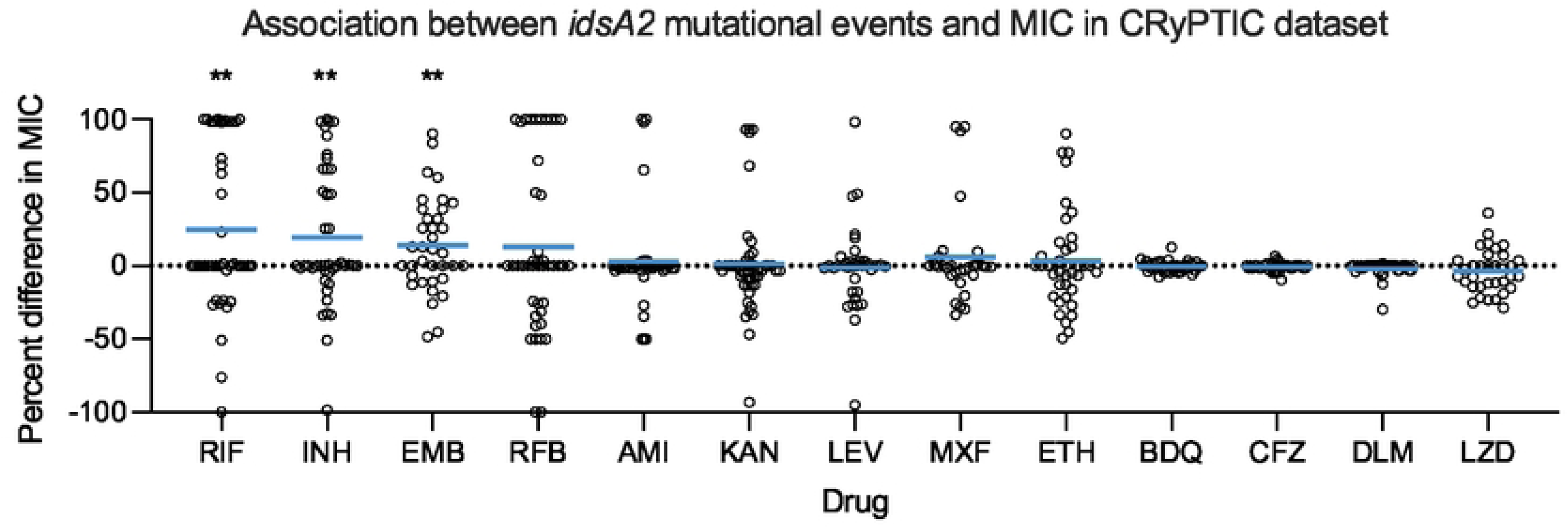
*idsA2* mutations are associated with drug resistance in clinical strains. Percent changes in minimum inhibitory concentration (MIC) of 13 drugs associated with *idsA2* mutational events in Lineage 4 data from the CRyPTIC consortium. Each dot represents the MIC difference attributed to an *idsA2* mutational event relative to its nearest neighbor strains, normalized using the maximum possible difference in MIC for the drug. The blue line indicates the average MIC difference across all *idsA2* mutational events. A permutation test was performed to assess significance of the average difference associated with *idsA2.* ** q<0.01.

### *idsA2* mutations alter sensitivity to first line antibiotics

We used oligo recombineering to introduce clinically identified *idsA2* variants into H37Rv, a Lineage 4 reference strain [19]. We chose two variants: one nonsense variant, S115X, representing an overt loss of function of IdsA2, and one missense variant, L204R, which was the most common *idsA2* mutation across Lineage 4 in our 55K dataset (Fig 1B). Because the causality of chromosomal single nucleotide variants cannot be established through complementation, we generated multiple independent clones for each allele and used whole genome sequencing to identify potential secondary site mutations. We also isolated recombineered clones without *idsA2* variants to use as wildtype controls.

We then assessed the impact of the engineered *idsA2* mutations on Mtb fitness in the presence and absence of drug. In standard axenic culture conditions, *idsA2* mutant strains had a minor, although statistically significant, growth defect relative to wildtype strains (S5 Fig). The strains’ antibiotic sensitivity was assessed for four first- and second-line antibiotics: ethambutol, isoniazid, rifampicin, and ofloxacin, quantitating the drug concentration at which 90% of bacterial growth was inhibited (IC90). Mutations in *idsA2* caused a roughly 2-fold increase in IC90 of ethambutol, a more modest increase in IC90 of isoniazid, as well as a small, but consistent decrease, in IC90 of rifampicin (Figs 3 and S5). These data align with findings from drug screening experiments with transposon-insertion and CRISPRi knockdown libraries [20,21]. Mutations in *idsA2* did not increase drug tolerance to any of the tested agents as measured by time-kill assays (S6 Fig).

**Fig 3:**
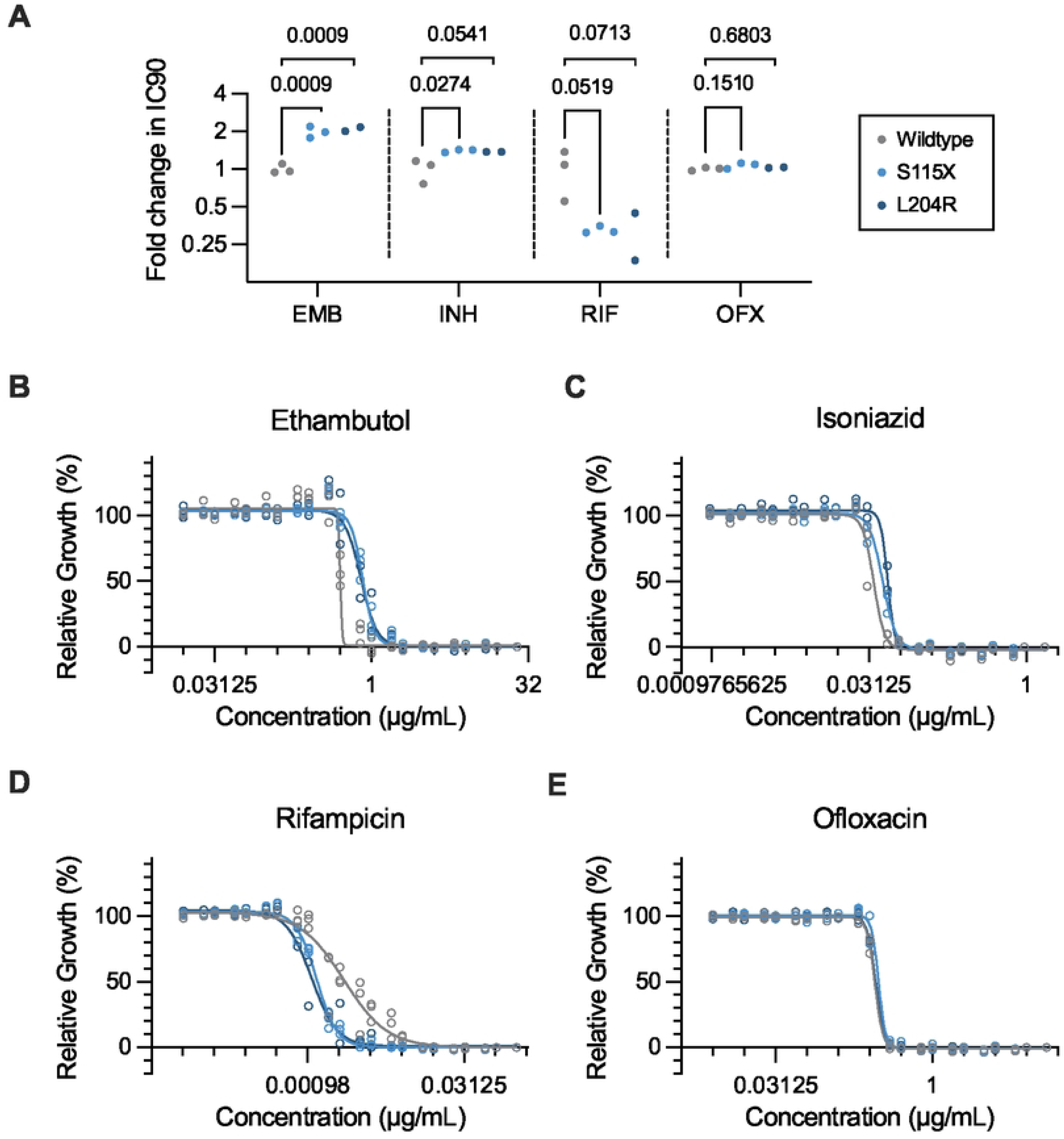
Isogenic *idsA2* mutants have altered sensitivity to first-line antibiotics. (A) Fold change in IC90 of wildtype and *idsA2* mutant strains in four tuberculosis antibiotics: ethambutol (EMB), isoniazid (INH), rifampicin (RIF), and ofloxacin (OFX), as measured by AlamarBlue reduction assay.

IC90 was calculated from nonlinear regression curves for each colony replicate from two experiments. Differences between each mutant and wildtype were tested by performing ordinary one-way ANOVA with Dunnett’s multiple comparison test on IC90 values (S5 Fig). (B-E) Nonlinear regression curves of growth in varying concentrations of antibiotics. Points come from two independent experiments, and each represents the mean of two (wildtype) to three technical replicates for each colony replicate.

### IdsA2 loss of function alters isoprenoid synthesis

To increase our confidence in the relationship between *idsA2* variants and these subtle but reproducible antibiotic effects, we next sought to define the metabolic consequences of *idsA2* disruption. Previous *in vitro* and structural analyses showed that purified IdsA2 catalyzes short-chain isoprenoid synthesis, subsequently allowing the synthesis of essential components such as cell wall precursors and menaquinone. Specifically, IdsA2 mediates the conversion of dimethylallyl pyrophosphate (DMAPP) through the intermediate molecule geranyl pyrophosphate (*E*-C_10_), to farnesyl pyrophosphate (C_15_) (Fig 4A) [11,22]. It is also capable of producing the downstream molecule geranylgeranyl pyrophosphate (*E*-C_20_) [23]. At each of these steps however, previous work suggests there is functional redundancy of IdsA2 with other prenyl diphosphate synthases such as GrcC2, IdsA1 and IdsB [23–25]. It was not clear to what extent these enzymes would compensate for IdsA2 disruption.

**Fig 4:**
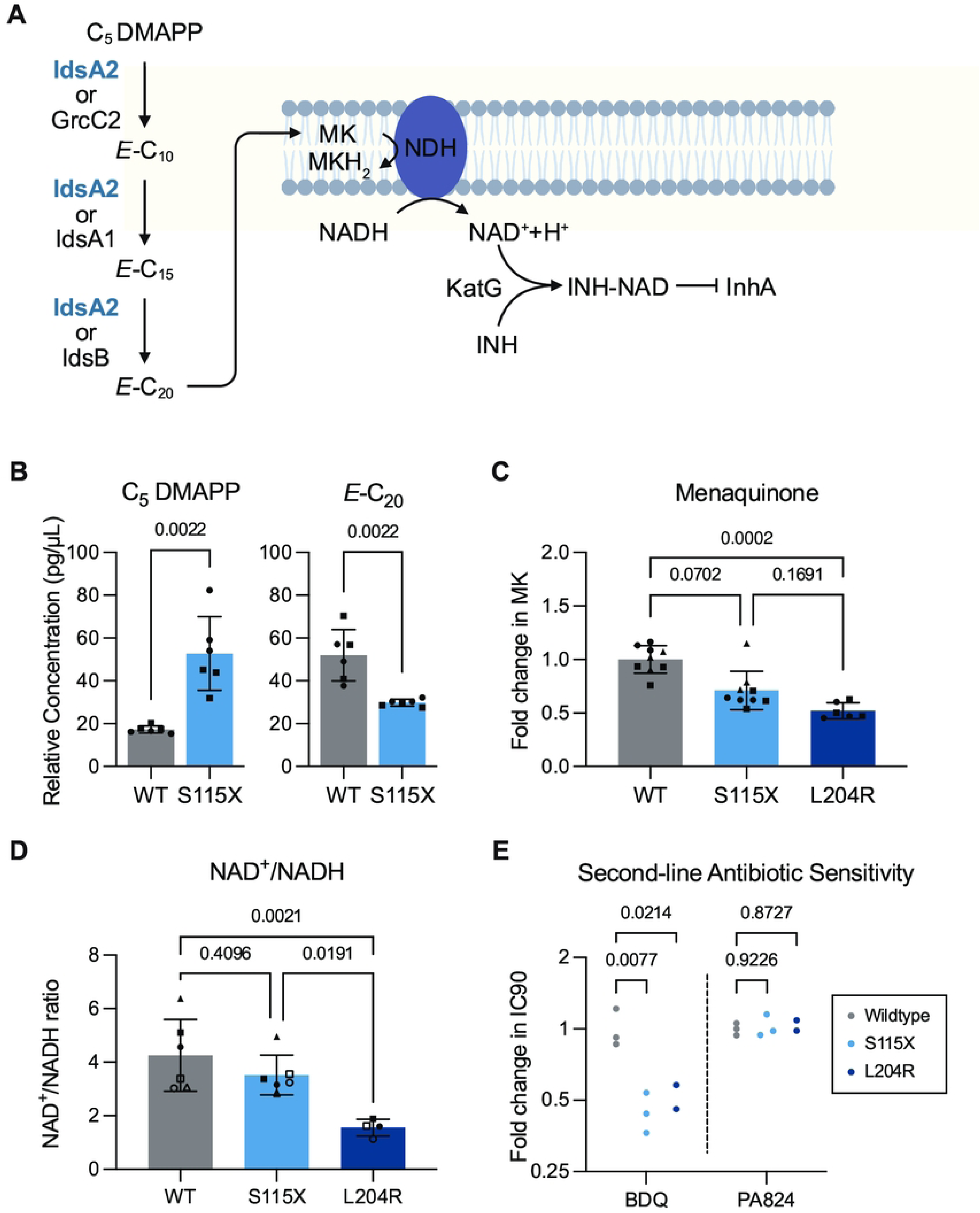
IdsA2 functionality effects isoprenoid synthesis and bacterial energetics. (A) Model of isoprenoid synthesis feeding into menaquinone (MK) production and the electron transport chain [11,26]. Made in BioRender. (B) C_5_ (DMAPP) and *E-*C_20_ concentrations in metabolite from wildtype and S115X mutant Mtb as measured by LCMS against standards added quantitatively. Dots are a combination of two colony replicates and three culture replicates. Different shapes indicate different colony replicates.

Differences between S115X and wildtype were measured by Mann-Whitney test for each molecule separately. (B) Total menaquinone as measured by LCMS. Dots are a combination of three colony replicates and three culture replicates. Different shapes indicate different colony replicates. Differences in menaquinone levels determined by Kruskal-Wallis test with Dunn’s multiple comparisons test. (C) Ratio of NAD^+^ to NADH determined by fluorometric assay. Data combines two independent experiments.

Differences in NAD^+^/NADH ratio determined by ordinary one-way ANOVA with Tukey’s multiple comparison test. (D) Fold change in IC90 of wildtype and *idsA2* mutant strains in bedaquline (BDQ) and pretomanid (PA-824) as measured by AlamarBlue reduction assay. IC90 was calculated from nonlinear regression curves for each colony replicate from one representative experiment for BDQ. IC90 was calculated from nonlinear regression curves for each colony replicate from two experiments for PA-824.

Differences between each mutant and wildtype were tested by performing ordinary one-way ANOVA with Dunnett’s multiple comparison test on IC90 values (S9 Fig).

To directly assess how *idsA2* disruption impacts this pathway, we sought to quantify the steady-state abundance of the predicted substrates and products of IdsA2 in the cell. To this end, we collected whole-cell lysate from our *idsA2* nonsense mutant strain (S115X) and compared to the wildtype strain using liquid chromatography mass spectrometry (LCMS). In the mutant strain, DMAPP concentrations were significantly increased and *E*-C_20_ concentrations were significantly decreased compared to the wildtype strain (Fig 4B). *E*-C_10_ and C_15_ were not detected, presumably due to their previously described rapid turnover which prevents accumulation as detectable pools [11,27]. These data show that loss of IdsA2 activity alters the short-chain isoprenoid synthesis pathway.

Since *E*-C_20_ is the precursor to menaquinone, we next used targeted lipid LCMS with collisional mass spectrometry to assess the levels of menaquinone in *idsA2* mutant and wildtype strains [28]. We found that total menaquinone levels in the missense mutant strain were significantly decreased compared to the wildtype strain, with the nonsense mutant trending in the same direction (Figs 4C and S7-8). Since menaquinone and NAD redox are coupled by NADH dehydrogenase activity [26,29,30], we then used a fluorescence assay to measure NAD^+^ and NADH concentrations and found that the ratio of NAD^+^ to NADH similarly trended down in both *idsA2* mutant strains (Figs 4D and S8). In previous experimental and genomic studies, decreases in NAD^+^/NADH ratios have been implicated in isoniazid resistance [31–35]. Therefore, the observed alteration in NAD^+^/NADH redox in the *idsA2* mutant strains represents a plausible mechanism for the small increase in the IC90 of isoniazid (Fig 3).

Isoniazid activity is impacted by, but does not directly target, bacterial energetics. However, newer WHO-recommended second-line antibiotics such as bedaquiline and pretomanid do. Therefore, we asked whether *idsA2* mutations also impact the sensitivity to these drugs. Interestingly, *idsA2* mutant strains were significantly more sensitive to bedaquiline, demonstrating the potential for collateral sensitivity caused by these mutations (Figs 4E and S9).

### *idsA2* mutants have increased levels of cell-wall precursors

We next asked whether the synthesis of cell wall precursors is also impacted by *idsA2* mutation. On this arm of the isoprenoid synthesis pathway, geranyl pyrophosphate (*E-*C_10_) is converted by Rv1086 into the *Z*-isomer of farnesyl pyrophosphate (*Z*-C_15_) which is then directly converted into decaprenyl phosphate, a crucial carrier lipid for synthesis of multiple constituents of the mycobacterial cell wall (Fig 5A) [36–38]. Directed analysis of the total abundance of decaprenyl phosphate species in *idsA2* mutant and wildtype strains found that steady-state levels of decaprenyl phosphate species were significantly increased in both the *idsA2* nonsense and missense mutant strains (Fig 5B). This data suggests a straightforward metabolic rerouting whereby IdsA2 impairment causes increased *E*-C_10_ which is converted into *Z*-C_15_ – presumably by Rv1086 – which is in turn used for synthesis of decaprenyl phosphate [36,37].

**Fig 5:**
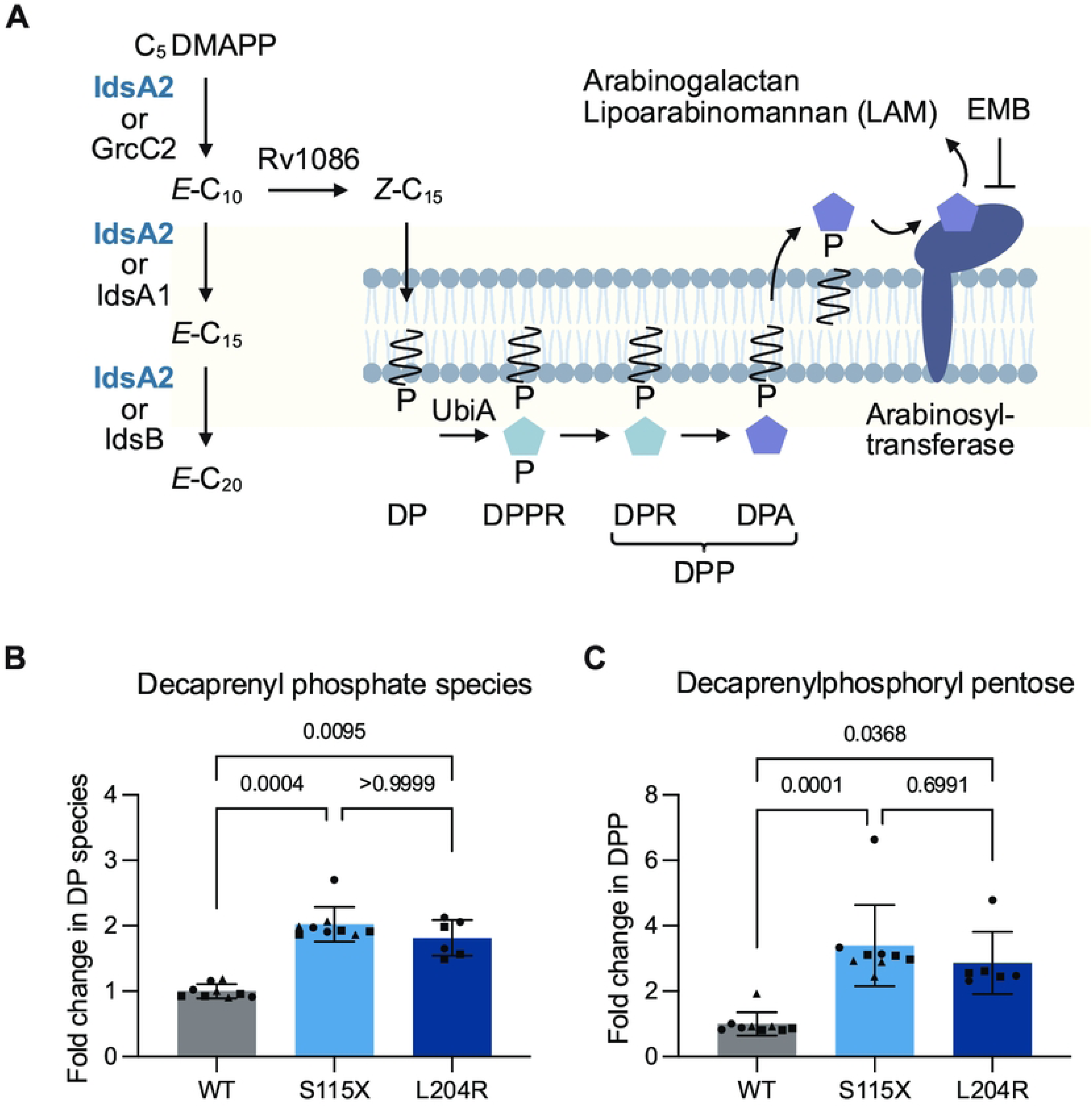
*idsA2* mutants have increased synthesis of DPA. (A) Model of isoprenoid synthesis feeding into arabinogalactan and LAM synthesis pathway [11,39]. Made in BioRender. (B-C) Total decaprenyl phosphate (DP) species and decaprenylphosphoryl pentose (DPP) as measured by LCMS. Dots are a combination of three colony replicates and three culture replicates. Different shapes indicate different colony replicates. Differences in DPP levels determined by Kruskal-Wallis test with Dunn’s multiple comparisons test.

Notably, one decaprenyl phosphate species with significantly increased levels in the *idsA2* mutant strains was decaprenylphosphoryl pentose (DPP), a combined measure of decaprenylphosphoryl arabinose (DPA) and decaprenylphosphoryl ribose (DPR) (Figs 5C and S10). Previous work has found that mutation or upregulation of the decaprenylphosphoryl-5-phosphoribose (DPPR) synthase, encoded by *ubiA*, also increased both DPA levels and ethambutol MIC [40–42]. Mechanistically, it was proposed that since DPA is the carrier molecule for arabinose [43,44], increasing the quantity of DPA allows arabinose to outcompete ethambutol for binding to its drug target, the arabinosyltransferase EmbB [42,45,46]. Our combined genomic, functional, and metabolic data support a model in which *idsA2* variants similarly increase the ethambutol MIC through increased production of DPA.

### *idsA2* mutations have multiplicative effects with *embB* mutations

Since mutation of EmbB, the arabinosyl transferase targeted by ethambutol, is the most common ethambutol resistance mechanism [47], we next sought to determine if *idsA2* mutations interact with mutations in *embB*. Using the Lineage 4 strains in the 55K dataset, we performed phylogenetic reconstruction to assess whether *embB* mutations occurred before or after *idsA2* mutational events. Excluding nodes in which ordering could not be determined, 90% of the time, *embB* mutations arose before *idsA2* mutations (Fig 6A). *EmbB* mutations arose before predicted-deleterious *idsA2* mutations 97% of the time (S11 Fig). This argues that *idsA2* variants are not primarily steppingstone mutations for acquisition of *embB* variants and instead suggests that they confer an additional advantage on top of existing ethambutol resistance.

**Fig 6:**
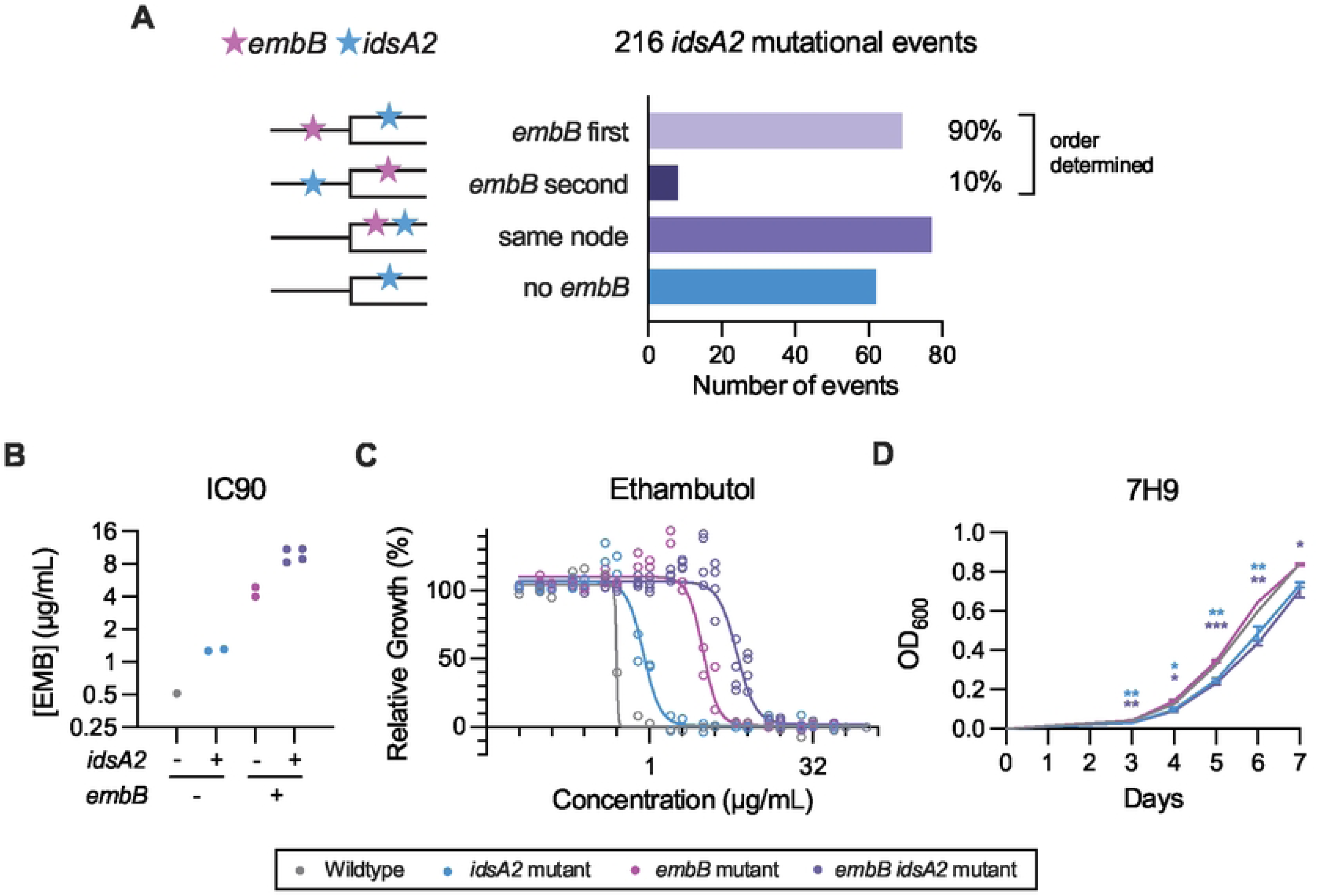
*idsA2* confers multiplicative drug resistance with *embB* mutations. (A) Ordering of 216 *idsA2* mutational events with *embB* mutations within Lineage 4 of the 55K dataset. (B) IC90 values calculated from nonlinear regression curves for each colony replicate from two experiments. (C) Nonlinear regression curves of growth in varying concentrations of ethambutol for each strain group.

Points come from two independent experiments, and each represents the mean of two (*idsA2* wildtype) to three technical replicates for each colony replicate. (D) Growth curves of wildtype, *idsA2* mutant, *embB* mutant, and combined *idsA2* and *embB* mutant strains in standard 7H9 media. Differences between each mutant and wildtype were tested by performing ordinary one-way ANOVA with Dunnett’s multiple comparison test for each day. * p<0.05, ** p<0.01, *** p<0.001.

To experimentally test the effects of *idsA2* variants in combination with *embB* mutations, we generated strains with mutations in both *embB* and *idsA2*. We plated *idsA2* wildtype and mutant strains on ethambutol to isolate independent clones with the ethambutol resistance-associated *embB* Q497R mutation in each strain background [47]. We then assessed the effects of the combined mutations on ethambutol resistance. In the absence of additional ethambutol-resistance mutations, *idsA2* mutations increased ethambutol IC90 by 0.5 µg/mL, resulting in an IC90 of 1 µg/mL (Fig 3 and 6B-C). In the presence of *embB* Q497R strain, however, *idsA2* mutations increased ethambutol IC90 by 4 µg/mL, resulting in an IC90 of 8 µg/mL (Figs 6B-C). An ethambutol MIC of 8 µg/mL is sufficient for a strain to be considered clinically resistant [48]. Thus, *idsA2* mutations can combine with *embB* mutations to elevate ethambutol resistance to clinically relevant concentrations.

An alternative explanation for *embB* and *idsA2* variant ordering could be that *idsA2* variants compensate for a loss of fitness due to *embB* mutation. However, the growth defect caused by the *idsA2* variants was maintained in the presence of the *embB* mutation, while *embB* Q497R itself does not alter replicative fitness in axenic culture (Fig 6D). Together, these data demonstrate that *idsA2* mutations augment ethambutol resistance.

### Presence of *idsA2* mutations can increase specificity of ethambutol susceptibility testing

There is a need to improve genetics-based ethambutol resistance diagnostics. Currently, the only gene with mutations associated with ethambutol resistance according to the WHO is *embB,* and there remain gaps in both the sensitivity and specificity of WHO-verified *embB* mutations for detecting phenotypic ethambutol resistance [47,49]. Knowing the effect of *idsA2* mutations on ethambutol resistance, we asked if there is clinical utility in identifying *idsA2* mutations for genotypic ethambutol susceptibility testing.

Returning to the CRyPTIC dataset, we first assessed the sensitivity and specificity of predicting ethambutol resistance using only WHO-verified ethambutol resistance mutations in *embB*. Since lineage typing would not typically occur at diagnosis, we used all strains regardless of lineage for this analysis. *EmbB-*based diagnosis of ethambutol resistance – here defining resistant strains as having an ethambutol MIC ≥4 µg/mL and sensitive strains as having MIC of ≤2 µg/mL – had a sensitivity of 85% and a specificity of 95% (Fig 7A). Since *idsA2* mutations often arise after *embB* mutations, inclusion of *idsA2* variants was not expected to increase sensitivity. However, specificity is also an important metric for diagnostics, as a high false positive rate can lead to unnecessary escalation to more toxic drug regimens. We hypothesized that checking for the existence of an *idsA2* mutation in an *embB* mutant strain may improve the specificity of detecting ethambutol resistance. Indeed, the specificity of using the combined presence of a WHO-verified *embB* mutation and an *idsA2* mutation for detection of ethambutol resistance was 99.93%, or a less than 1% false positive rate.

**Fig 7:**
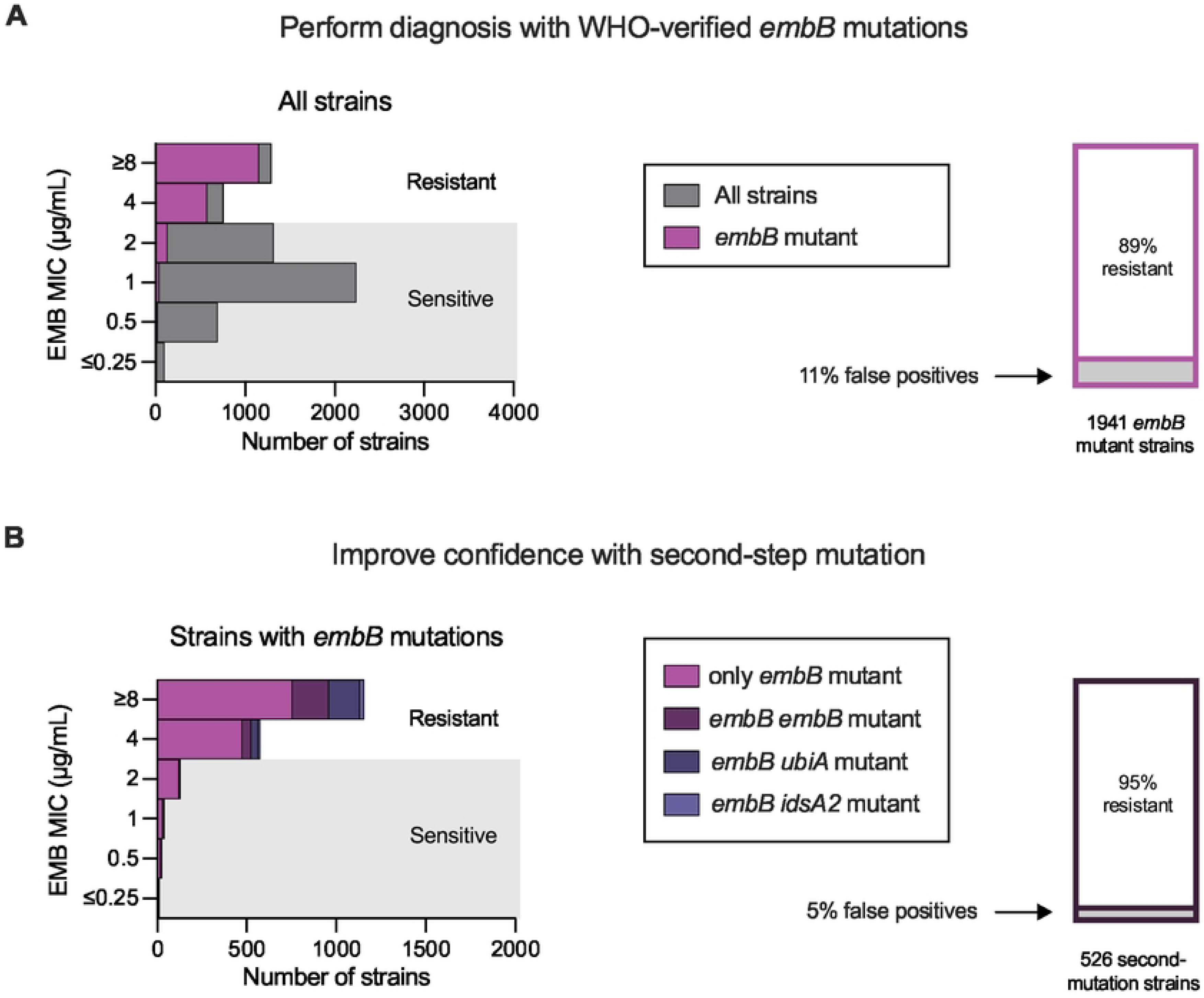
Second-step ethambutol resistance mutations improve diagnostic specificity. (A, left) Distribution of ethambutol MICs across all strains in the CRyPTIC dataset. 11% of strains with WHO-verified *embB* mutations have an MIC ≤2 µg/mL meaning they would be false positives for diagnosing ethambutol resistance. (B) Distribution of ethambutol MICs across WHO-verified *embB* mutant strains. 5% of strains with a WHO-verified *embB* mutation and a second step mutation in *idsA2, ubiA,* or *embB*, have an MIC ≤2 µg/mL meaning they would be false positives for diagnosing ethambutol resistance.

Since *idsA2* mutations were present in <2% of *embB* mutant strains in our dataset, we next asked if we could improve the frequency at which this two-step approach would be effective by detecting mutations in other genes which represent the second step to ethambutol resistance. Mutations in *ubiA* and the presence of more than one mutation in *embB* have both been associated with higher ethambutol MICs [42,50]. Repeating our analysis with these variants, we found that in both cases the specificity was approximately 99.7% for prediction of ethambutol resistance, again indicating a less than 1% false positive rate. Furthermore, approximately 27% of *embB* mutant strains had additional mutations in at least one of these three genes (Fig 7B).

Therefore, 27% of the time, the specificity in diagnosing ethambutol resistance can be improved to over 99% by identifying secondary-site mutations in strains with WHO-verified ethambutol resistance variants.

## Discussion

### The pleiotropic effects of *idsA2* mutation on drug susceptibility

In this study, we demonstrate that *idsA2* is under selection in a lineage-specific manner and, in Lineage 4 Mtb strains, loss of IdsA2 enzymatic function is associated with first-line drug resistance. IdsA2 loss of function remodels the isoprenoid synthesis pathway, resulting in wide-ranging impacts on essential processes from cell wall synthesis to bacterial energetics. This pathway remodeling also enables its most clinically relevant effect, amplifying ethambutol resistance after *embB* mutation.

The pleiotropic effects of *idsA2* mutations may also impact the efficacy of newer antimycobacterial agents. Encouragingly, we found that *idsA2* mutant strains were more susceptible to bedaquiline. However, efficacy of other antibiotics may suffer. Quabodepistat, a drug which targets DprE1, one of the two essential enzymes involved in the conversion of DPR to DPA, is currently undergoing Phase 3 clinical trials for treatment of drug-resistant TB in a combination therapy [51]. Although not identified in studies of Quabodepistat, studies of other DprE1 inhibitors have identified *ubiA* mutations as causing low-level resistance [52,53].

Therefore, it is plausible that increased synthesis of DPR could outcompete DprE1 inhibitors from binding to their targets, and that this new drug class could therefore be less effective against strains with *ubiA* or *idsA2* mutations. Additionally, *idsA2* mutations spontaneously developed in the presence of a new thieno[2,3-b]pyridin-6(7H)-one derivative [54]. Together, this illustrates the importance of testing new antimycobacterial compounds on strains with existing drug resistance.

### Validating IdsA2 functionality in whole-cell assays

*In vitro* studies of isoprenoid synthesis enzymes were essential to our understanding of this pathway; however, they incompletely account for the complex whole-cell environment.

Here, our analysis of the steady-state abundance of isoprene-based molecules in whole-cell lysate from wildtype and *idsA2* mutant bacteria allowed us to infer the role of IdsA2 in the context of functionally redundant enzymes and validate previous findings [11]. Aligning with structural evidence, the simplest explanation for the significant accumulation of DMAPP in the IdsA2 loss-of-function strain is that IdsA2 uses DMAPP as a substrate in the whole-cell context [22]. Furthermore, although we could not directly detect high turnover intermediates *E*-C_10_ or *E*-C_15_, we can infer that IdsA2 uses *E*-C_10_ as a substrate, as decreased competition for *E*-C_10_ would allow for metabolic rerouting and the observed increase in decaprenyl phosphate-species in the *idsA2* mutant strains. Finally, previous work has found that IdsA2 can synthesize *E*-C_20_ *in vitro* [11,24]. We found that *E*-C_20_ levels were decreased in our IdsA2 loss-of-function strain, however it is unclear whether this is due to loss of IdsA2 as a redundant *E*-C_15_ to *E*-C_20_ synthesizing enzyme or due to decreased *E*-C_15_ substrate availability. Overall, our work confirms previous findings that IdsA2 is a DMAPP/*E*-C_10_ to *E*-C_10_/*E*-C_15_ synthase, with a potential role in *E*-C_20_ synthesis.

### Selection on the isoprenoid pathway

Ethambutol is of moderate potency as an antimycobacterial agent, and it is perhaps not surprising that the cell has discovered multiple ways to evade ethambutol pressure. As discussed above, although mutating the drug target *embB* is the most common way to acquire ethambutol resistance, *ubiA* mutations also contribute to ethambutol resistance by increasing DPA levels [42]. In this work, we provide evidence that *idsA2* mutations contribute to ethambutol resistance through the same mechanism, and we propose that they may not be the only other mutations to do so. Interestingly, published work assessing signals of selection in the Mtb genome across the four major lineages found modest signals of diversifying selection in *grcC2* and *idsB –* genes encoding enzymes with functional redundancy to IdsA2, acting at the DMAPP to *E*-C_10_ and *E*-C_15_ to *E*-C_20_ steps respectively [23,24] – in Lineage 2 [3]. Further work will be required to determine if mutations in these genes or others phenotypically replicate *idsA2* mutants. Further work is also required to understand the reason for these lineage specific signals of selection. One possibility is that there may be a preference for acquisition of a mutation in one lineage over another due to an epistatic interaction with a lineage-defining mutation. For *idsA2,* a variant private to Lineage 4 in *rv1086*, which encodes the protein which converts *E*-C_10_ to *Z*-C_15_, is a plausible explanation for the Lineage 4 specific *idsA2* selection [36].

### The challenge of genomically predicting ethambutol resistance

This work shows how both lineage-specific selection and genetic epistasis can obfuscate the importance of a variant in drug resistance. These issues likely contribute to the challenge of identifying genetic markers of ethambutol resistance, where there remain gaps in both the sensitivity and specificity of known variants to detect phenotypic ethambutol susceptibility [47,55]. These gaps are due in part to the continuous nature of ethambutol resistance. The field has traditionally sought out single variants to explain and detect high-level drug resistance. This approach is well suited for drugs such as isoniazid where the MICs have a bimodal distribution due to the large and consistent jump in MIC caused by single variants. However, ethambutol MICs distribute unimodally with a long tail skewing up towards resistance [48]. This and other work show that this is not only due to *embB* mutations conferring varying levels of resistance, but due to the stepwise increases of MIC on top of *embB* variants caused by mutations in *idsA2*, *ubiA*, and additional *embB* mutations [42,50]. We show that by leveraging an understanding of combinatorial ethambutol resistance, the specificity of predicting ethambutol resistance can be increased. As the field improves modeling for predicting phenotypic drug resistance, lineage and sub-lineage effects as well as multi-step evolution should be considered.

## Conclusion

In this work, we show that *idsA2* variants contribute to first-line drug resistance through the pleiotropic effects of metabolic remodeling. Most notably, *idsA2* mutations have a multiplicative effect with *embB* variants on ethambutol resistance. This adds *idsA2* to a group of second-step ethambutol resistance mutations including mutations in *ubiA* and second-site mutations in *embB.* Detection of these second-step mutations can increase the specificity of *embB* mutation*-*based ethambutol resistance prediction. These results not only help close the gap in our genotypic understanding of phenotypic drug susceptibility in Mtb but also emphasize the importance of considering the genetic interactions that might create lineage specific or combinatorial drug phenotypes that might be relevant to patient care.

## Methods

### Genomic analyses

The two datasets used in this study are referred to as the 55K dataset and the CRyPTIC dataset. We used a table of *pN/pS* values, calculated from the 55K dataset for previously published work [3], but excluding reversions (S1 Table). Variant calling and tree building for the 55K dataset was performed for a previous study [3]. Resulting phylogenetic trees, ancestral reconstruction, and variant calling were deposited to Mendeley (DOI: 10.17632/h9dd6hsvc2.1). The CRyPTIC Consortium has generously made their dataset publicly available. We downloaded the CRyPTIC_reuse_20231208.csv and used the ENA_RUN information to pull whole genome sequences from the Short Read Archive (SRA) and perform variant calling using the same pipeline as used for the 55K dataset with H37Rv as a reference [3]. Resulting phylogenetic trees, ancestral reconstruction, and variant calling were deposited to Mendeley (DOI: 10.17632/cy4dy352kj.1). Included in this pipeline, annotations and effects for each variant were called with *snpeff* and *sift4g,* [17,56]. Whenever we refer to variants in a specific gene, we are referring to any variants annotated as being within the gene and meeting the following criteria: disruptive inframe deletion or insertion, frameshift, missense, and stop variants. When we refer specifically to predicted-deleterious *idsA2* mutations, those were variants annotated as frameshift or stop variants by *snpeff* or deleterious missense variants by *sift4g*. *tbprofiler* was used to assign lineages for each strain; only Lineage 4 strains were used for all analyses with the exception of Fig 1A and Fig 7 [57,58]. Code used for the following analyses is available in the associated Github (https://github.com/abigailmfrey/IdsA2_Paper_Code). All analyses were performed using the Harvard FAS Research Computing (FASRC) cluster.

### Variant plotting

Variants within the *idsA2* coding region (Gene ID Rv2173) and meeting the variant requirements described above were pulled from the annotated event calls for the whole dataset of interest. Lineage 4 strains with these variants were identified. The number of strains with each variant was calculated, and then variants were grouped by HGVS number per annotation type.

Missense variants were designated as deleterious or tolerated by *sift4g* [17]. Disruptive inframe insertions and deletions were combined, and any combination of annotations including a frameshift was counted as a strain with a frameshift variant. The results were plotted in lollipop form across the gene or in pie chart form to demonstrate abundance of variant type using GraphPad Prism 10.

### Loss of function prediction analysis

To compare loss of function signals across the genome, the rate of frameshift and nonsense mutations per kilobase per gene was calculated using the 55K dataset. Frameshift and stop variants were pulled from the event calling, removing variants with a miss rate of 5% or higher. Variants were also removed if they fell in a region of the genome deemed unreliable for variant calling or had more than one reversion [59]. Genes must have had at least 80% of the gene callable and have at least 100 bases. The number of frameshifts and stop variants per gene was then divided by the number of kilobases in the callable region of the gene resulting in events per kilobase.

### Variant association with MIC in CRyPTIC dataset

To assess the association of *idsA2* variants with MIC changes across the 13 drugs from the CRyPTIC dataset, we sought to determine the difference in MIC between strains resulting from *idsA2* mutational events and their nearest neighbors. Only MIC values defined as high quality for the drug being analyzed were used. Due to differences in plate set-up, the extreme values were adjusted to reflect only the values available on both plates [48]. For example, UKMYC5 has an ethambutol range of 0.06-8 mg/L with minimum MIC values labeled as <=0.06 mg/L and maximum >8 mg/L. UKMYC6 has an ethambutol range of 0.25-32 mg/L with minimum MIC values labeled as <=0.25 mg/L and maximum >32 mg/L. For our analysis, all values less than or equal to 0.25 (<=0.06, 0.12, <=0.25, 0.25) from both plates were labeled as 0.25 while all values greater than or equal to 8 (8, >8, 16, 32, >32) were labeled as 8. Variant calling, tree building, annotation, and variant settings are described above. The CRyPTIC tree was pruned using *dendropy* to only include Lineage 4 strains with high quality MIC data for the drug of interest [60]. We then calculated the difference in MIC between each target and sibling set, otherwise referred to as the nearest neighbor(s). We then identified the nodes with *idsA2* mutational events per our standard settings. We averaged the calculated difference between the event node and its nearest neighbors across all events to acquire the associated change in MIC for *idsA2* mutational events. We then performed a permutation test, averaging the calculated differences across the same number of random nodes as *idsA2* events in that tree, 10,000 times. Significance was determined by how many times the test returned an average difference greater than the *idsA2* events. Multiple testing was corrected by two-stage linear step-up procedure of Benjamini, Krieger and Yekutieli in Graphpad Prism 10 with a 1% false discovery rate.

### *Mycobacterium tuberculosis* isogenic mutant strain construction

*IdsA2* point mutants were constructed using oligo-mediated recombineering as previously described [2,19]. Specifically, a strain of H37Rv was transformed with two plasmids, pKM427 (a chromosomal site L5 integrating vector carrying a zeocin resistance cassette and a hygromycin cassette that is inactivated by a premature stop codon) and pKM402 (an episomal vector carrying the phage recT recombinase gene under an ATc inducible promoter). The resulting strain was grown to early log phase, induced with ATc, then transformed with two single-strand DNA oligos. One oligo confers hygromycin resistance and the other oligo carries the *idsA2* mutation of interest (S115X or L204R). The transformants were recovered for 2 days in 7H9 (7H9 salts with 0.2% glycerol, 10% OADC, 0.05% Tween-80) then plated on 7H10 (7H10 media with 0.5% glycerol, 10% OADC, 0.05% Tween-80) containing hygromycin (50 μg/mL). Colonies were screened for *idsA2* mutations by Sanger sequencing. Three colonies were retained for each mutant of interest and three hygromycin-resistant colonies with no *idsA2* mutations were retained for use as wildtype controls. All strains were expanded and re-streaked onto solid media with hygromycin and colonies which had spontaneously cured the pKM402 plasmid were saved. The resulting strains were whole genome sequenced (200 Mbp Illumina DNA sequencing, SeqCenter). Whole genome sequencing reads have been deposited onto the SRA (BioProject ID PRJNA1356913). Variants relative to H37Rv were identified using *breseq* [61]. Variants found in all strains, in masked regions, or found to be inconsistently called within our dataset were not included in S4 Table [59]. There were no additional variants shared between clone replicates (S4 Table). However, a single base pair deletion in *papA5*, a gene involved in PDIM biosynthesis, was found in one L204R clone. Due to the loss of PDIM enhancing bacterial growth rate, this clone was dropped from the study [62]. Unless otherwise indicated, experiments were conducted using three independently derived clones of wildtype and S115X strains, and two independently derived clones of the L204R strain.

### Growth curves

Mid-log phase cultures grown in 7H9 at 37°C shaking were diluted to an OD_600_ 0.001 in duplicate in 15 mL 7H9. OD_600_ was measured on days 3-7. Differences between each mutant and wildtype were tested by performing ordinary one-way ANOVA with Dunnett’s multiple comparison test. Two independent growth curves were performed.

### Antibiotic resistance measurement

Antibiotic resistance against four commonly used anti-tuberculosis drugs (EMB, INH, RIF, and OFX) and two second line antibiotics (BDQ and PA824) was assessed using AlamarBlue assays as previously described with the following modifications [2,8]. Strains were grown to mid-log phase 7H9 and diluted to a final OD_600_ of 0.0015 in in 200 µL 7H9 media containing antibiotics at indicated concentrations in a 96-well plate. After five days of incubation at 37°C with constant agitation, AlamarBlue reagent was added at 10% of the culture volume.

The plates were then incubated at 37°C for one day before measuring absorbance. The amount of AlamarBlue reduction was estimated by subtracting background OD_600_ from OD_570_. The percentage of bacterial growth was calculated on a row-by-row basis by subtracting the relative absorbance measure of the highest drug concentration well from the relative absorbance measure of the no-drug control well. Strains were plated in duplicate (WT) or triplicate (mutant) rows.

The antibiotic concentration at which growth is inhibited by 90% (IC90) for each clone replicate was approximated by performing nonlinear regression on data from two independent experiments performed with different starting concentrations using GraphPad Prism 10. For bedaquiline, calculations were performed for the two experiments independently and one representative experiment is shown. The three wildtype clones’ IC90s were averaged and used to calculate the IC90 fold-change for each clone of each strain per drug. P-values represent differences between each mutant and wildtype IC90 as tested by performing ordinary one-way ANOVA with Dunnett’s multiple comparison test. Nonlinear regression curves are shown for data where replicate rows were averaged, and regression was performed for all clone replicates together for visual clarity.

### Antibiotic tolerance measurement

Antibiotic tolerance against four commonly used anti-tuberculosis drugs (EMB, INH, RIF, and OFX) was assessed using a time-kill assay as previously described with the following modifications [2]. Strains were grown to mid-log phase in 7H9 and diluted to a final OD_600_ of 0.05 in in 10 mL 7H9 media in a 50 mL inkwell. Antibiotics were added to final concentrations of 100x the calculated MIC of our wildtype strain (55, 3, 0.6, and 34 µg/mL respectively). The cultures were then incubated at 37°C with constant agitation and sampled at day 1, day 3, and day 5. On each day, a 1 mL aliquot of each antibiotic-supplemented culture was taken, and the cells were washed and resuspended in 1mL 7H9 without antibiotic. After washing off the antibiotics, a series of dilutions determined by pilot experiments were prepared in a 96-well plate using fresh 7H9 media. 5μL of each dilution was spotted on a 7H10 agar plate with no antibiotics. Colony enumeration was performed at 24 days after plating.

### Metabolite extraction

Two clone replicates of wildtype and S115X were grown in technical triplicate to mid-log phase in 7H9. 10 mL of culture was spun down at 3000 rpm for 10 minutes at 4°C then the pellet was resuspended in 1 mL of chilled 2:2:1 acetonitrile:methanol:water. Bacteria were then lysed by bead beating 0.5 min x 6 resting on ice in between cycles. Tubes were then spun down at 14,000 rpm for 10 minutes at 4 °C. Supernatant was removed and filtered twice for removal from the BL3.

### LC-MS/MS analysis of isoprenoid pyrophosphates

Isoprenoid pyrophosphates were analyzed using an LC–MS/MS system equipped with a Phenomenex Luna C18 column (150 × 2 mm) by the Harvard Center for Mass Spectrometry. Mobile phase A consisted of 10 mM ammonium bicarbonate, and mobile phase B was methanol. Standards were purchased in 1 mg/mL concentrations in methanol from Sigma (D4287, G6772, F6892, G6025) and run at known concentrations to generate a standard curve. 10 µL aliquots of standards and samples were injected for each run. The flow rate was maintained at 0.3 mL/min. The LC gradient program was as follows: 0 min, 0% B; 6 min, 99% B; 10 min, 99% B; 10.1 min, 0% B; 13 min, 0% B. Detection was performed in negative ionization mode using multiple reaction monitoring (MRM). MRM transitions and corresponding parameters for each analyte were as follows:

- **DMAPP (C_5_)**: m/z 245 → 78.9 (fragmentor 80 V, CE 17 V, cell voltage 4 V) and m/z 245 → 158.8 (fragmentor 80 V, CE 9 V, cell voltage 4 V)
- **GPP (*E*-C_10_)**: m/z 313.1 → 78.9 (fragmentor 90 V, CE 21 V, cell voltage 5 V) and m/z 313.1 → 294.9 (fragmentor 90 V, CE 4 V, cell voltage 4 V)
- **FPP (C_15_)**: m/z 381.1 → 78.9 (fragmentor 110 V, CE 21 V, cell voltage 13 V) and m/z 381.1 → 158.8 (fragmentor 110 V, CE 13 V, cell voltage 4 V)
- **GGPP (*E*-C_20_)**: m/z 449.2 → 79 (fragmentor 140 V, CE 33 V, cell voltage 4 V) and m/z 449.2 → 158.8 (fragmentor 140 V, CE 13 V, cell voltage 4 V)

Source parameters were set as follows: gas temperature, 350 °C; gas flow, 12 L/min; nebulizer pressure, 35 psi; sheath gas temperature, 400 °C; sheath gas flow, 12 L/min. Concentrations of DMAPP and GGPP were normalized based on protein concentration of the sample as determined by microBCA. Difference between S115X and wildtype were measured by Mann Whitney test for each molecule separately.

### Lipid extraction

Lipid extraction was done as previously described [63]. Cultures were grown in 7H9 to mid-log phase. They were then spun down and washed with detergent free media then diluted to OD_600_ = 0.0075 in 45 mL 7H9 without detergent in triplicate. Cultures were then grown to OD_600_ = 0.55-0.7. Cultures were spun at 4000 rpm for 10 minutes and then washed twice with 10 mL Optima LCMS grade water. Pellet was resuspended in 1 mL methanol. Resuspended solution was then added to a glass vial containing 3 mL methanol and 2 mL chloroform creating a final ratio 1:2 chloroform:methanol (C:M). Samples were left for 30 minutes at room temperature to sterilize before removal from the BL3. Samples were then incubated on a rotator for 1 hour at room temperature. Cells were spun down at 3000 rpm for 10 minutes at room temperature and supernatant was transferred to a new glass vial. Pellet was resuspended in 1:1 C:M and incubated for 1 hour on rotator at room temperature. Spin was repeated and supernatant was collected and added to previous supernatant. Supernatant was then evaporated using Genevac. Dried lipids were resuspended in 4 mL 1:1 C:M. Lipid suspension was sonicated at 60 sonics/minute for 3 minutes, shaken throughout. Samples were again spun down and then supernatant was transferred to a pre-weighed amber glass vial. Lipids were dried using Genevac. Vials were then weighed again to acquire final total lipid mass. All weighing was performed using a Mettler Toledo XP205 analytical balance with an EN 8 SLC deionizing power supply. Lipids were resuspended to 1 mg/mL 1:1 C:M and stored at -20°C.

### Lipid LCMS and collisional mass spectrometry

Lipid HPLC-MS and collisional induced dissociation mass spectrometry (CID-MS) were carried out using a reversed phase method [64]. Lipid preparations at 1 mg/mL in 1:1 C:M were evaporated under nitrogen and dissolved to the same volume in solvent A (95:5 methanol:water with 2 mM ammonium formate). 10 μg (10 μL) lipid was injected per sample, using an Agilent 1260 infinity HPLC with an Agilent Poroshell 120 EC-C18 column (1.9 μm particle size, 3.0 x 50 mm, 699675-302), connected to an Agilent 6530 electrospray ionization quantitative time-of-flight (ESI-QTOF) mass spectrometer. A flow rate of 0.15 mL/minute was used, with the following ratios of solvent A to solvent B: 0-4 minutes, 100% A; 4-10 minutes, gradient to 100% B; 10-15 minutes, 100% B; 15-20 minutes, gradient to 100% A; followed by a 5 minute postrun with 100% A. Solvent B was 90:10:0.1 1-propanol:cyclohexane:water with 3 mM ammonium formate. Ionization was maintained at 325°C gas temperature, 8 L/minute flow rate of drying gas, 35 PSI nebulizer pressure, and 3500 V. Spectra were collected at 2 spectra/second from 100 to 3200 m/z, in both the positive and negative ion modes, over separate runs. CID-MS was performed on the same instrument with the same conditions, detecting a 40-3000 m/z range with collision energy of 20-50 V, using a medium (4 m/z) mass window, starting from the chosen ion m/z rounded down to the nearest whole number minus 0.3 (e.g. for ion m/z 909.6378, CID-MS was carried out from 908.7-912.7 m/z).

### Lipid measurement

All analysis of lipid mass spectral data was performed using Agilent Masshunter Qualitative Analysis software. Menaquinones were detected in the positive mode, and decaprenyl species in the negative mode, by searching for exact m/z matches. The single ionized adduct with the highest intensity was chosen for each lipid. Individual chromatographic peaks were visualized and structurally validated by CID-MS and integrated in a sample-blinded manner, in profile mode, with a 10 ppm m/z window and no smoothing. Selected bar plots show fold change values calculated for each strain’s clone and technical replicates relative to an average value taken across all wildtype samples. There is significantly more variability within technical replicates than between clone replicates as shown in S10 Fig, allowing us to plot technical and clone replicates together. Total decaprenyl phosphate species values were calculated by summing the intensities of each decaprenyl phosphate species we could detect: decaprenyl phosphate, decaprenylphosphoryl pentose (putatively a combination of ribose and arabinose) and decaprenylphosphoryl hexose (putatively a combination of glucose and mannose). Similarly, total menaquinone levels were calculated by summing intensities of menaquinones and dihydromenaquinones 8 and 9. P-values were determined by Kruskal-Wallis test with Dunn’s multiple comparisons test.

### NAD/NADH measurements

NAD/NADH measurements were performed with the Fluorescent NAD/NADH Detection Kit (Cell Technology). Briefly, cultures were grown to mid-log phase in 7H9, normalized by OD_600_, and 1 mL or 3 mL of culture were taken for NAD or NADH measurements respectively. Samples were pelleted by centrifugation, washed twice with PBS, and resuspended in extraction buffer. Lysis buffer and bead beating lysed the bacteria and samples were incubated at 60°C for 1.5 hours. On ice, reaction and the opposite extraction buffer were added. The plate setup was performed as described with half the volume and after 1 hour incubation was read at 550 nm excitation and 595 nm emission.

### Ordering analysis

Variant events meeting our criteria occurring in *idsA2* in Lineage 4 strains from the 55K dataset were pulled from event calling. We then asked which strains following this event had *embB* variants. First, if none of the strains had any *embB* mutation, the event was labeled as having only *idsA2.* Next, if a certain *embB* variant was called as a “miss” in all strains under the *idsA2* event, indicating poor variant calling, the variant was thrown out for this event. Next, if all strains were found to have the same *embB* variant, we then went back to the parent node and asked if all strains under that node have the same *embB* variant indicating that the *embB* variant happened at the parent node or earlier. If all strains under the parent node had the *embB* variant, the event was called as *embB* happening first. If there was any other combination of calls within the parent strains (no variant, variant, and/or misses) the *embB* event is assumed to occur at the same node as the *idsA2* event. Finally, if only some strains resulting from the *idsA2* event had an *embB* variant, this is called as the *embB* event(s) coming second.

### Acquisition of ethambutol resistance mutations

A representative clone for each of our strains of interest (*idsA2* WT, S115X, L204R) were grown on ethambutol at 4 or 8 µg/mL. Several colonies were then picked, grown up in 7H9, and gDNA was extracted. WGS was performed (200 Mbp Illumina DNA sequencing, SeqCenter) and reads were mapped to H37Rv using *breseq* [61]. Two strains from each background with *embB* Q497R mutations were used for further experiments.

### Sensitivity and specificity of ethambutol resistance predictions

Analysis was performed using strains of all lineages in the CRyPTIC dataset. Variant calling criteria and MIC handling is described above. Strains were considered sensitive to ethambutol if they had an MIC at ≤2 µg/mL, and resistant if they had an MIC of ≥4 µg/mL. WHO-verified *embB* mutations are those defined as associated with resistance to ethambutol in the second edition of the drug resistance catalog [47]. After identifying the presence of WHO-verified *embB* mutations, our variant calling criteria were then used to determine whether each strain had mutations in *idsA2, ubiA,* or any additional *embB* mutation. Sensitivity and specificity were calculated for each variant condition of interest.

## Acknowledgements

We would like to thank Michael Chao for his critical reading of this manuscript.

## Supporting Information

**S1 Fig: Distribution of *pN/pS* ratios for L1, L2, and L3.** Histograms of *pN/pS* ratios for all genes in Mtb from the Lineages 1, 2, and 3 from the 55K dataset. Median values indicated by dotted line are 0.64, 0.75, and 0.67 respectively. *idsA2 pN/pS* values indicated are 0.61, 0.81 and 0.87 respectively.

**S2 Fig: *IdsA2* variants in the CRyPTIC dataset.** (A) The number of strains per codon from Lineage 4 of the CRyPTIC dataset with disruptive in-frame deletions or insertions, frameshift, deleterious or tolerated missense, or nonsense mutations in *idsA2*. (B) Pie chart of total percentages of each variant type among Lineage 4 strains in the CRyPTIC dataset.

**S3 Fig: Association between *idsA2* mutational event and MIC change in the CRyPTIC dataset.** Difference in MIC of 13 drugs associated with *idsA2* mutational events in Lineage 4 data from the CRyPTIC consortium. Each dot represents the MIC difference attributed to an *idsA2* mutational event relative to its nearest neighbor strains. The blue line is the mean MIC difference across all *idsA2* mutational events. A permutation test was performed to assess the significance of the mean difference associated with *idsA2* and a multiple test correction was performed with an FDR of 1%.

**S4 Fig: Association between predicted-deleterious *idsA2* mutational event and MIC change in the CRyPTIC dataset.** Difference in MIC of 13 drugs associated with predicted-deleterious *idsA2* mutational events in Lineage 4 data from the CRyPTIC consortium. Each dot represents the MIC difference attributed to an *idsA2* mutational event relative to its nearest neighbor strains. The blue line is the mean MIC difference across all *idsA2* mutational events. A permutation test was performed to assess the significance of the mean difference associated with *idsA2* and a multiple test correction was performed with an FDR of 1%.

**S5 Fig: Growth curve and IC90 statistics.** (A) Growth curves of wildtype and *idsA2* mutant strains in standard 7H9 media. Differences between each mutant and wildtype were tested by performing ordinary one-way ANOVA with Dunnett’s multiple comparison test for each day. * p<0.05, ** p<0.01, *** p<0.001 **** p<0.0001. (B) Plots of IC90 values for each strain separated by drug: ethambutol (EMB), isoniazid (INH), rifampicin (RIF), and ofloxacin (OFX). Differences between each mutant and wildtype were tested by performing ordinary one-way ANOVA with Dunnett’s multiple comparison test.

**S6 Fig: Kill curve assay.** (A) Minimum duration to 90% cell death (MDK90) for wildtype and *idsA2* mutant strains in four tuberculosis antibiotics: ethambutol (EMB), isoniazid (INH), ifampicin (RIF), and ofloxacin (OFX). The concentration of each drug was 55, 3, 0.6, and 34 µg/mL. MDK90 was calculated using a semilog curve equation determined using the line segment through which Fraction Surviving = 0.01 fell. Differences between each mutant and wildtype for each drug were tested by ordinary two-way ANOVA with Dunnett’s multiple comparison test. (B-F) Fraction of surviving bacteria over time calculated from colony enumeration.

**S7 Fig: Representative chromatograms and collision spectra for menaquinone species.** Red boxes on extracted ion chromatograms (EICs) indicate peaks with lipid identity confirmed by CID-MS via the highlighted fragments. For each lipid, displayed prenyl stereochemistry and position of saturated groups is assumed from literature patterns and may differ across chromatographic peaks. Blue diamonds in spectra indicate the lowest targeted m/z value for CID -MS.

**S8 Fig: Supporting data for menaquinone and redox analysis.** (A) Intensity measured in arbitrary units (counts) for measured menaquinone species. (B) The concentrations of NAD^+^ and NADH as measured by fluorometric assay. Data combines two independent experiments.

**S9 Fig: IC90 statistics and nonlinear regression curves for second-line antibiotics.** (A,C) Plots of IC90 values for each strain separated by drug: bedaquiline (BDQ) and pretomanid (PA-824). Differences between each mutant and wildtype were tested by performing ordinary one-way ANOVA with Dunnett’s multiple comparison test. (B,D) Nonlinear regression curves of growth in varying concentrations of antibiotics. Points each represents the mean of two (wildtype) to three technical replicates for each colony replicate. For BDQ, the plot is representative of two experiments. For PA-824, points come from two independent experiments.

**S10 Fig: Supporting data for polyprenyl analysis.** (A-C) Representative chromatograms and collision spectra for decaprenyl phospholipids. Red boxes on EICs indicate peaks with lipid identity confirmed by CID-MS via the highlighted fragments. For each lipid, displayed prenyl stereochemistry and position of saturated groups is assumed from literature patterns and may differ across chromatographic peaks. Blue diamonds in spectra indicate the lowest targeted m/z value for CID -MS. (D) Intensity measured in arbitrary units (counts) for measured decaprenyl phosphate species. (E) Fold change in DPP levels as measured by LCMS where each bar is a clone, and dots are technical replicates. Different shapes indicate different colony replicates corresponding with Fig 5C. Differences in DPP levels determined by Kruskal-Wallis test with Dunn’s multiple comparisons test.

**S11 Fig: Ordering of predicted-deleterious *idsA2* mutational events.** Ordering of 158 predicted-deleterious *idsA2* mutational events within Lineage 4 of the 55K dataset.

**S1 Table: *pN/pS* calculations without reversions.** *pN/pS* values were generated into a histogram for Fig 1A and S1.

**S2 Table: Lineage 4 defining variants.**

**S3 Table: Number of nonsense and frameshift events per gene per lineage.**

**S4 Table: Strain list.** Includes strain names and variants identified by WGS after filtering.

**S1 Data: Data underlying plots.** Excludes Fig1A and S1 (S1 Table) and Fig 1D (S3 Table).

## Notes

### Competing Interest Statement

SMF receives compensation as a non-executive director of Oxford Nanopore Technologies (ONT). No ONT sequencing was performed in this study.

